# Variants in *Lrrk2* and *Snca* deficiency do not alter the course of primary encephalitis due to neurotropic reovirus T3D in newborn mice

**DOI:** 10.1101/2024.09.28.615578

**Authors:** Michaela O. Lunn, Christopher Rousso, Julianna J. Tomlinson, Earl G. Brown, Michael G. Schlossmacher

**Author notes:** **Correspondence and requests for materials:** These should be addressed to J.J.T or M.G.S. Lead contacts (Julianna J. Tomlinson, Earl G. Brown, Michael G. Schlossmacher).

## Abstract

Variants of the leucine-rich repeat kinase-2 (*LRRK2*) and α-synuclein (*SNCA*) genes are associated with Parkinson’s disease risk. We previously demonstrated that two *Lrrk2* knock-in variants as well as *Snca* expression alter survival rates from combined pneumonitis and encephalitis following intranasal inoculation of newborn mice with a double-stranded RNA virus: respiratory-enteric-orphan virus, serotype-3 strain Dearing (reovirus T3D). Here, we examined whether outcomes of direct inoculation of the brain by reovirus T3D, which invariably causes lethal encephalitis within 15 days, would also be modified by variants in *Lrrk2* and *Snca*. When we inoculated newborn mice intracerebrally with 5×10^2^ plaque-forming units of reovirus T3D, we found that, when compared to wild-type littermates, Lrrk2 p.G2019S mutant mice and kinase-dead p.D1994S mutant animals showed the same time-to-death intervals post-infection, revealed no sex difference, and had similar viral titres in the brain. Furthermore, the reduction or absence of endogenous α-synuclein also did not alter the course of encephalitis in parallel studies. These outcomes are in contrast to those following the intranasal inoculation paradigm of newborn mice, in which Lrrk2 and wild-type α-synuclein were both protective in infection outcomes. Together, these findings suggest that the Parkinson’s disease-linked *Lrrk2* and *Snca* genes contribute predominantly to systemic, innate responses by the host following reovirus T3D exposure.

## Introduction

Parkinson’s disease (PD) is one of the most prevalent neurodegenerative disorders, characterized by the progressive loss of dopaminergic neurons in the Substantia nigra pars compacta leading to a typical movement disorder. While the exact etiology of PD remains elusive, the pathogenesis likely requires a multifaceted interplay between genetic and environmental factors. Initiation of PD pathology may begin outside of the brain, such as in the nose and/or gut, two sites that are in direct contact with the external environment (Kalia and Lang, 2015; Braak et al., 2003). Consistent with a theory that PD could result from environmental - possibly microbial - triggers, epidemiological data suggest that the incidence rate of prior infections with several viruses, including hepatitis C virus, herpes simplex virus, and influenza A virus are significantly correlated with elevated PD risk (Leta et al., 2022). There is increasing interest in elucidating the role of environmental triggers and the relative contribution of peripheral versus CNS-specific host responses, in the initiation and progression of PD pathologies.

*LRRK2* and *SNCA* are two loci linked to altered risk regarding the development of PD. We and others have shown in mice that both genes are involved in the response to viral and bacterial infections *in vivo* (Shutinoski et al., 2019; Härtlova et al., 2018; Liu et al., 2017; Zhang et al., 2015; Alam et al., 2022; Monogue et al., 2022; Tomlinson et al., 2017; Beatman et al., 2016; Chen et al., 2016). Furthermore, we found that the PD-linked p.G2019S mutation in Lrrk2 conferred a hyper-inflammatory phenotype in infected mice, leading to better control of both *Salmonella typhimurium* growth and reovirus T3D titres following intravenous and intranasal inoculation, respectively. However, when the infection reached the brain in the case of reovirus T3D, Lrrk2 p.G2019S knock-in mice had worse health outcomes indicated by increased mortality, which was also associated with a significant female sex bias. Contrarily, Lrrk2 kinase-dead p.D1994S neonates were relatively protected from intranasal infection by reovirus T3D, with increased survival rates when compared to their wild-type littermates.

Both *Lrrk2* and *Snca* are expressed in the brain and the periphery. The respective, relative contribution of their involvement in host defences in the periphery versus the CNS to respond to reovirus T3D infection-induced encephalitis remains unknown. Here we sought to address this by assessing *Lrrk2*- and *Snca*-dependent outcomes following a direct-brain inoculation of murine pups with reovirus T3D to compare outcomes with our previous studies using an intranasal-inoculation paradigm with the same virus (Shutinoski et al., 2019; Tomlinson et al., 2017). Intriguingly, neither *Lrrk2* variants nor *Snca* expression altered acute disease outcomes, including time-to-death intervals and viral titres following direct-brain inoculation, suggesting that both proteins may play a critical role in peripheral organs to protect brain health following infection by this RNA virus.

## Methods

### Mouse Colonies

Animal experiments were performed under the guidelines by Canadian Councils on Animal Care and approved by the Ethics Board of the Animal Care Committee at the University of Ottawa. Experiments involving viral infections were done in a containment level 2 biohazard animal facility. Lrrk2 p.G2019S mice and Lrrk2 p.D1994S mice are described in Herzig et. al. (2011). α-Synuclein knockout (*Snca*^*-/-*^) mice were first described by Cabin et. al. (2002) and were kindly obtained from Dr. Matthew Farrer. All mice are on a C57Bl/6J background. Both female and male mice were used in this study and are indicated within each figure.

### Intracerebral Reovirus T3D Infection

From heterozygous breeding pairs, post-natal day 1 (p1) mouse pups were anesthetized under 3% isoflurane in oxygen. Using a 30-gauge 50μL fixed needle Hamilton syringe, 10μL of live, purified Respiratory Enteric Orphan Serotype 3 Dearing Virus (reovirus T3D) in PBS at a dose of 5×10^2^ PFU or 10μL PBS alone was injected into the left hemisphere of the pup brain (frontal lobe) (Passini and Wolfe, 2001) and returned to the home cage.

### Intracerebral Reovirus T3D Infection Survival Assay

Infected animals were assessed for survival, measured as time-to-death, using human endpoints. Mice were monitored twice daily (morning and evening) up to 14 days post inoculation (dpi).

### Viral Quantification in Tissue

A separate cohort of infected mice were euthanized at 8dpi via rapid decapitation for viral titre analyses in the brain and liver. Samples in PBS at a 3:1 ratio (μl PBS:mg tissue) for brains, or a 5:1 ratio for livers, were homogenized with metal beads using a Roche MagNA Lyser (Roche, Indianapolis, IN, USA) at maximum speed for 10 seconds. Samples underwent a freeze-thaw cycle in liquid nitrogen and were then centrifuged to collect the viral content in the supernatant for standard reovirus plaque assay analysis as described (Shutinoski et al., 2019). Briefly, tissue homogenates were used for serial dilutions and overlaid onto L929 cells in 6-well plates at 90-100% confluency. Infected cells were overlaid with 2% purified agar and 2x199 media in a 1:1 ratio. Plaques were identified after 7 days of incubation using 0.015% neutral red stain and manually counted. PFU was reported as plaques per gram of tissue.

### Statistical Analysis

All statistical analyses were performed using GraphPad Prism version 8 (GraphPad Software, San Diego, CA, USA, www.graphpad.com). Difference between groups in survival assays was assessed using the log-rank (Mantel-Cox) test. Differences between groups were assessed using one-way ANOVA with Tukey’s post-hoc tests. Data are demonstrated as mean ± SEM where applicable and as described in the figure legends. Data are displayed with *P* values represented as * *P*□<□0.05.

## Results

### Parkinson’s-linked Lrrk2 p.G2019S variant does not alter acute outcomes in a primary reovirus T3D-induced encephalitis model

The Lrrk2 p.G2019S variant was examined for its impact on acute outcomes in reovirus T3D-induced encephalitis following intracerebral inoculation. Lrrk2 p.G2019S mouse pups at postnatal day 1 were injected intracerebrally with reovirus T3D and either monitored for time-to-death or used for tissue collection and viral titre analyses (**Fig. 1a**). Mice carrying the p.G2019S allele (heterozygous or homozygous) had similar time-to-death intervals as did wild-type mice (*n* ≥ 7 mice per group) (**Fig. 1b, c**). Previously, we consistently observed a female sex bias with Lrrk2 in infectious disease states; therefore, we analyzed the data separately based on sex, but found no difference (**Fig. 1b, c**).

**Fig. 1:**
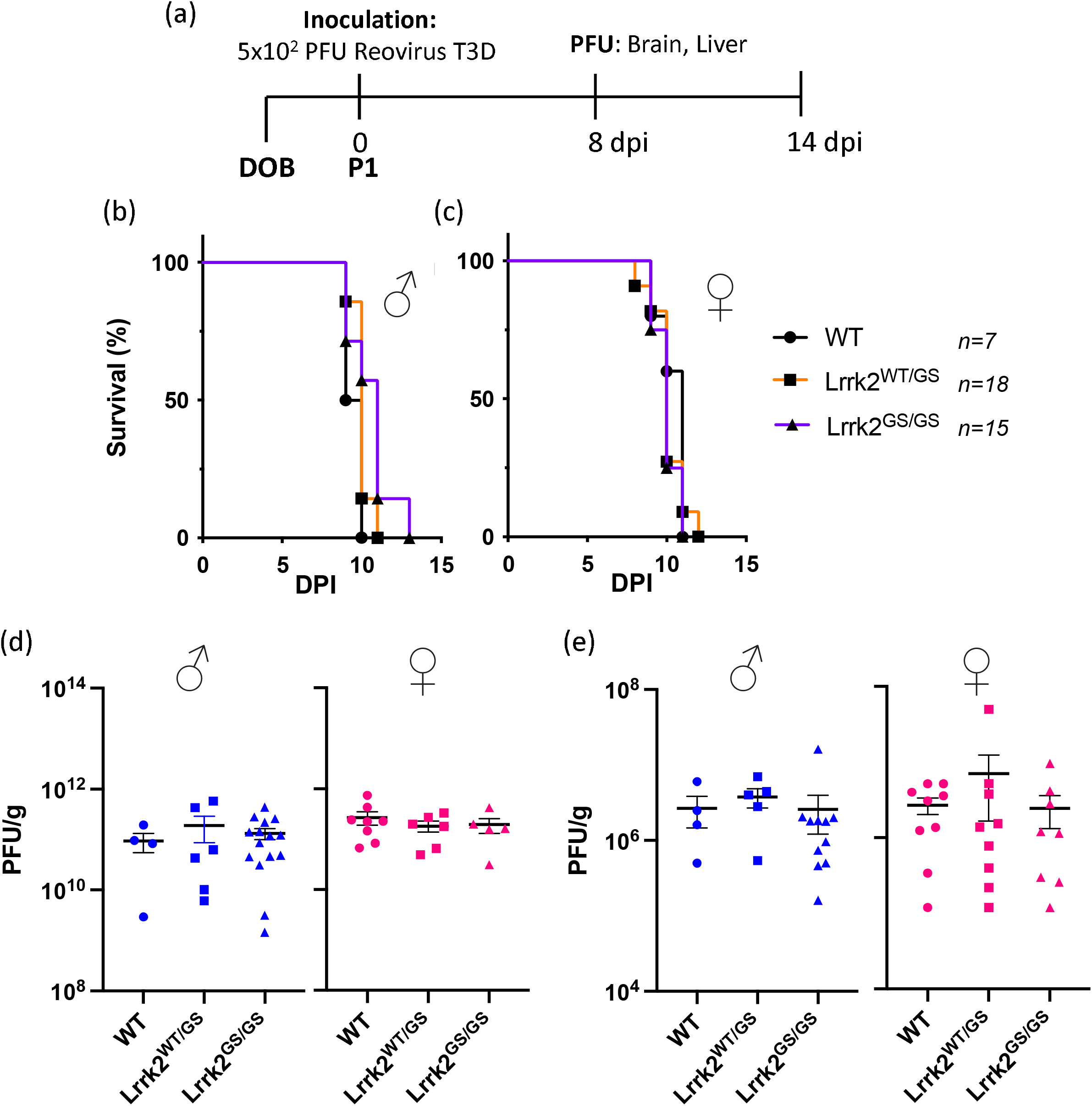
Expression of the Parkinson’s Disease-Linked Lrrk2 p.G2019S Variant Does Not Alter Outcomes of Acute Reovirus T3D-Induced Encephalitis When Inoculated Intracerebrally. Heterozygous Lrrk2 p.G2019S breeders were paired, and newborn mice were inoculated with a dose of 5×10^2^ PFU of reovirus T3D via direct brain injection into the left hemisphere and monitored for survival or sacrificed for tissue collection, as summarized in (**a**). Inoculated pups were monitored over 14 days for a pre-determined moribund state. Time-to-death is graphically displayed as percent survival against days post inoculation (DPI), separated based on sex (**b**). The *n* value indicates the sample size of sexes combined. In a second cohort, inoculated pups were sacrificed at 8dpi; brain and liver were collected and homogenized. Viral titres of the brain (**c**) and the liver (**d**) were measured via plaque assay. Data are represented in PFU/g, where each symbol represents one animal. Blue data points indicate male mice (*n* ≥ 4 per group) and pink data points indicate female mice (*n* ≥ 5 per group). Survival curves were statistically analyzed using Mantel-Cox (Log-Rank) tests; viral titres were analyzed using one-way ANOVA and Tukey post-hoc tests. No significant group differences were observed.

To assess viral infectivity and its replication in the brain, we quantified viral titres at 8 days post-infection as the latest timepoint before mice begin to succumb to the infection. Viral titres in the brain were similar among all genotypes, regardless of sex (**Fig. 1d**). We also investigated secondary dissemination of the pathogen from the brain by measuring viral titres in the liver at 8dpi and observed viral spread from the brain to the periphery. However, there were no differences in viral titres between wild-type and Lrrk2 p.G2019S expressing mice (**Fig. 1e**). Collectively, these findings indicated that the Lrrk2 p.G2019S hyper-kinase mutation did not alter the course of acute illness following intracerebral inoculation with reovirus T3D.

### Lrrk2 kinase-dead mice have an equal survival rate as heterozygous and wild-type mice following direct brain inoculation with reovirus T3D

We next investigated whether Lrrk2 kinase activity was required for the host’s defence against intracerebral inoculation with reovirus T3D; thus, we conducted the same experiments (as above) in kinase-dead Lrrk2 p.D1994S knock-in mice (Herzig et al., 2011). Unlike the survival benefit conferred by this mutation following intranasal inoculation (Shutinoski et al., 2019), there was no difference in time-to-death between wild-type and heterozygous and homozygous Lrrk2 p.D1994S mice (*n* ≥ 13 mice per group) following direct brain inoculation (**Fig. 2a**). Consistent with this, Lrrk2 p.D1994S mice had similar viral titres in the brain at 8dpi compared to wild-type and heterozygous mice (**Fig. 2b**). We observed viral dissemination to the liver at 8dpi in the Lrrk2 p.D1994S cohort, with relatively low but comparable viral titres across the three genotypes (**Fig. 2c**). Collectively, these findings suggested that neither p.D1994S nor p.G2019S allelic variants in Lrrk2 influenced the course of encephalitis after direct injection of reovirus T3D into the brain.

**Fig. 2:**
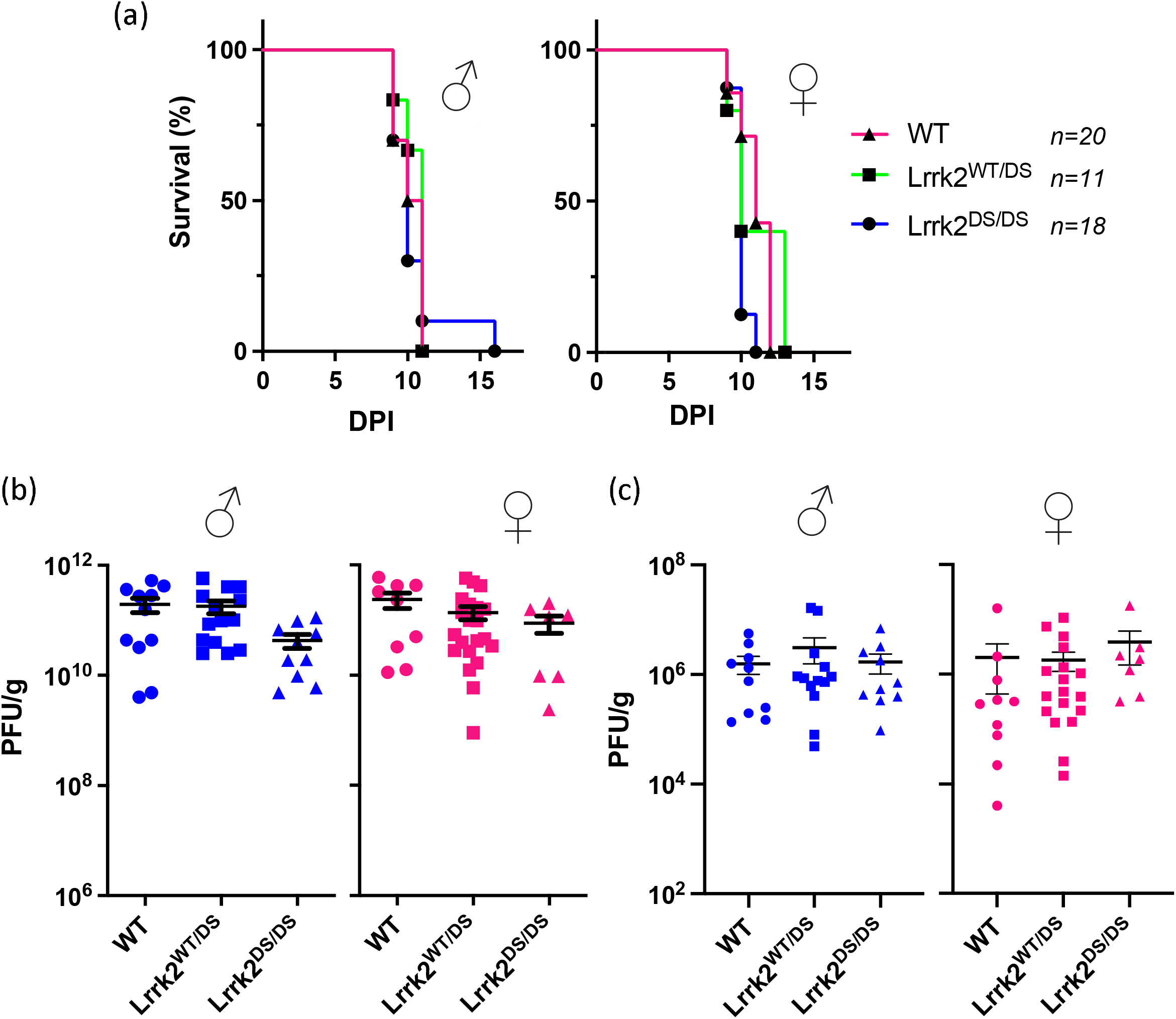
Expression of the Lrrk2 Kinase-Dead p.D1994S Mutant Does Not Alter Outcomes of Acute Reovirus T3D-Induced Encephalitis When Inoculated Intracerebrally. Heterozygous Lrrk2 p.D1994S breeders were paired, and newborn mice were inoculated with a dose of 5×10^2^ PFU of reovirus T3D via direct brain injection into the left hemisphere. Inoculated pups were monitored over 14 days for a pre-determined moribund state. Time-to-death is graphically displayed as percent survival against days post inoculation (DPI), separated based on sex (**a**). The *n* value indicates the sample size of sexes combined. In a second cohort, inoculated pups were sacrificed at 8dpi and brain and liver were collected and homogenized. Viral titres of the brain (**b**) and liver (**c**) were measured via plaque assay. Data are represented in PFU/g, where each symbol represents one animal. Blue data points indicate male mice (*n* ≥ 9 per group) and pink data points indicate female mice (*n* ≥ 7 per group). Survival curves were statistically analyzed using Mantel-Cox (Log-Rank) tests and viral titre data were statistically analyzed using one-way ANOVA and Tukey post-hoc tests. No significant group differences were observed.

### Endogenous α-synuclein does not protect against intracerebrally acquired reovirus-induced encephalitis

Akin to Lrrk2 and building on our previous discovery that *Snca* expression was protective against intranasal inoculation of pups with reovirus T3D (Tomlinson et al., 2017), we also investigated the antiviral properties of α-synuclein to protect against a primary encephalitis. We performed direct brain inoculation of reovirus T3D in *Snca*^-/-^, *Snca*^+/-^, and wild-type littermates. In contrast to intranasal, systemic administration of reovirus T3D, direct brain inoculation did not result in any genotypic differences in time-to-death, regardless of sex (*n* ≥ 10 mice per group) (**Fig. 3a**). Similarly, *Snca* expression also did not affect viral replication rates in the brain at 8dpi, regardless of sex (**Fig. 3b**).

**Fig. 3:**
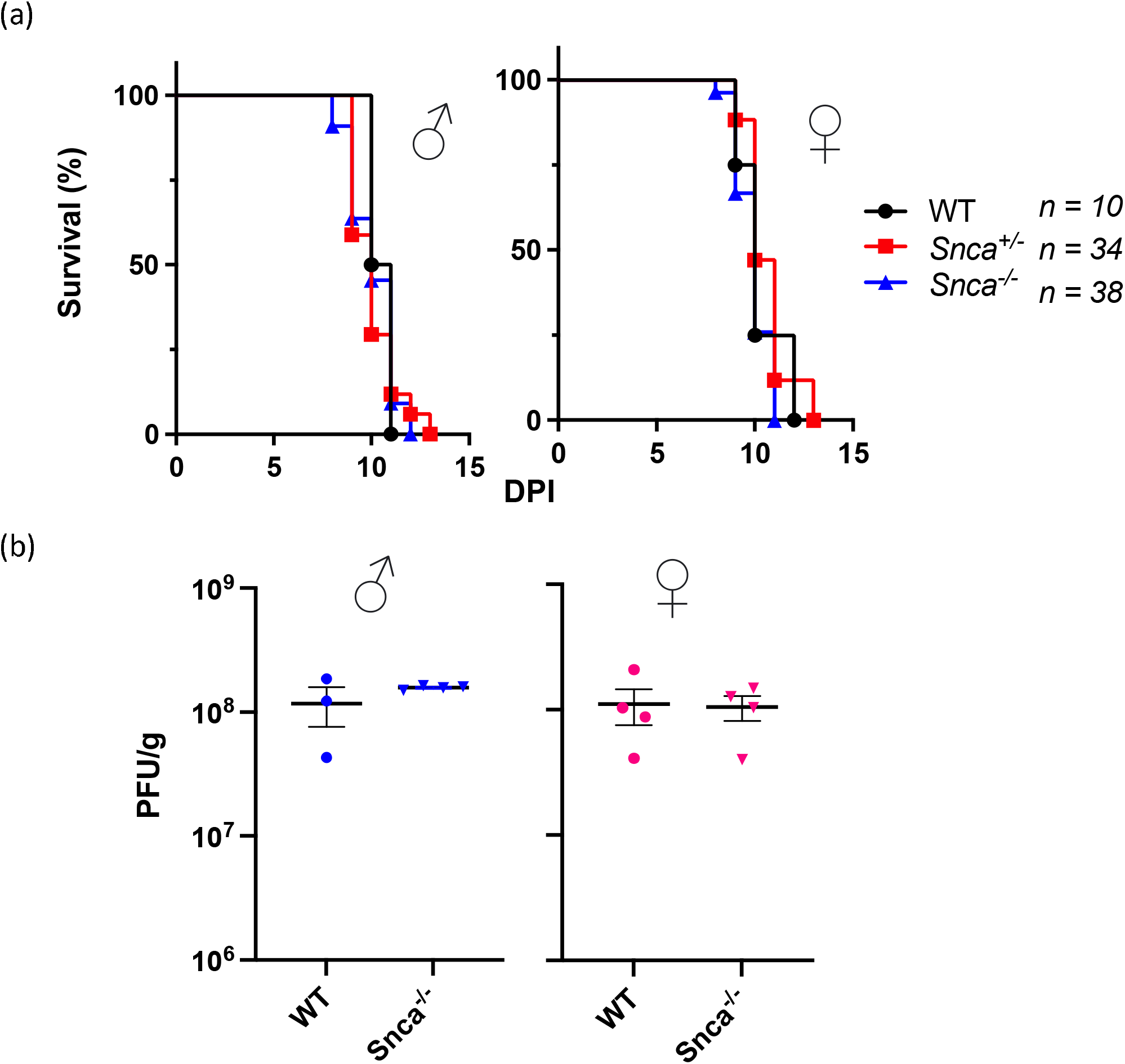
Expression of Murine *Snca* Gene Does Not Alter Outcomes of an Acute, Intracerebral Infection by Reovirus T3D-Induced Encephalitis. Newborn wild-type mice, *Snca* knockout (*Snca*^-/-^), heterozygous (*Snca*^+/-^), and wild-type (WT) pups were inoculated with a dose of 5×10^2^ PFU of reovirus T3D via direct brain injection into the left hemisphere. Inoculated pups were monitored over 14 days for a pre-determined moribund state. Time-to-death is graphically displayed as percent survival against days post inoculation (DPI), separated based on sex (**a**). The *n* value indicates the sample size of sexes combined. In a second cohort, inoculated pups were sacrificed at 8dpi and brains were collected for homogenization. Viral titres of the brain were measured via plaque assay (**b**). Data are represented in PFU/g, where each symbol represents one animal. Blue data points indicate male mice (*n* ≥ 3 per group) and pink data points indicate female mice (*n* ≥ 4 per group). Survival curves were statistically analyzed using Mantel-Cox (Log-Rank) tests and viral titre data were statistically analyzed using one-way ANOVA and Tukey post-hoc tests. No significant group differences were observed.

## Discussion

Host responses to microbial infections are pathogen- and tissue-dependent. We have previously shown that two PD-linked genes, *Lrrk2* and *Snca*, confer anti-microbial functions in mice. In the case of Lrrk2, we showed that this occurred in a pathogen-, tissue-, and sex-dependent manner. Specifically, while female Lrrk2 p.G2019S mice were better able to control reovirus T3D and *Salmonella typhimurium* titres in peripheral organs, in the case of intranasal inoculation of reovirus T3D, which subsequently infected the brain, mutant mice had greater disease severity with decreased survival due to encephalitis. Both of these genes are expressed within the CNS – including in neurons and microglia – as well as in the periphery. Immune responses that occur in the periphery have critical effects on brain health. In this current study, we sought to determine the relative contribution of Lrrk2 and α-synuclein function(s) in the immediate antiviral host response as it relates to survival outcomes from encephalitis following direct inoculation of the brain. Intranasal- and intracerebral inoculations of murine pups cause lethal encephalitis as a result of infection of the brain following a similar time course of disease but different routes (Gauvin et al. 2013).

In contrast to our expectations, in this study neither allelic variants in *Lrrk2* nor *Snca* expression itself impacted survival or viral replication following intracerebral inoculation. This outcome suggests to us that both genes can modulate peripheral host responses to acute reovirus T3D exposure, and that it is the peripheral involvement which conferred partial neuroprotection from otherwise lethal encephalitis when the infection had started outside the nervous system. When the virus was directly injected into the brain, these genes had no detectable impact on survival. Notably, both intranasal and intracerebral inoculation of reovirus T3D resulted in similar symptomatology and illness progression, with peak lethality occurring between 6-10dpi in both models.

Recent literature reports support the concept that Lrrk2 may influence pathogenic outcomes in the brain by acting in the periphery rather than directly in the brain. For example, Kozina et al. (2022) demonstrated not only that the peripheral administration of a bacterial antigen, lipopolysaccharide (LPS), could drive dopaminergic neuronal loss in transgenic Lrrk2 mutant mice (p.R1441G and p.G2019S), but also that it was dependent on the presence of peripheral monocytes; further, neutralization of peripheral IL-6 prevented the LPS-induced cell loss in transgenic, Lrrk2-mutant mice. Their findings support the hypothesis that allelic variants in *Lrrk2* may exert modulatory effects on immune cells and cytokines in the periphery, which in turn confer secondary, deleterious effects on dopaminergic neuronal health.

Akin to Lrrk2, albeit less-studied in this context, α-synuclein has been reported to have anti-microbial properties both *in vivo* and *in vitro*, including the protection of mice from viral and bacterial infections that began in the periphery (Tomlinson et al., 2017; Beatman et. al., 2016). Furthermore, α-synuclein has been shown to be involved in normal inflammatory responses to pathogens in the periphery (Alam et al., 2022). Increased *Snca* expression and post-translational modification have been reported downstream of microbial and inflammatory exposures (Peelaerts et al., 2023; Bantle et al., 2019; Jang et al., 2009), suggesting that the gene and protein may respond to environmental triggers as part of a physiological role in host responses; these findings could also link environmental exposures and ensuing host responses to PD-relevant changes in α-synuclein.

A key difference between intranasal and intracerebral inoculation models for reovirus T3D in suckling pups is that while the lethality of the infection following intranasal inoculation is titre-dependent, the median lethal dose of reovirus T3D delivered via direct inoculation of the brain is 10^1^ PFU (Virgin et al., 1988; Spriggs and Field, 1982). This represents a limitation of our study, in that the severity of the intracerebral inoculation method may be too fulminant to reveal time-to-death differences between genotypes in contrast to the intranasal reovirus T3D infection paradigm. However, previous literature showed that caspase-3 knockout mice had partial survival using the same intracerebral reovirus T3D infection model, suggesting that our model is sensitive enough to detect survival differences between specific genotypes (Beckham et al., 2011). To determine whether post-natal age at infection could affect the sensitivity of the assay (i.e., decreased lethality), we inoculated a cohort of mice at 3 days after birth, allowing for further brain development; nonetheless, time-to-death intervals were identical to those when animals had been infected on postnatal day 1, with again no differences seen between genotypes (**Supp. Fig. 1**).

A further limitation of this study is the use of neonatal pups, devoid of an adaptive immune system. While this model has allowed us to address a role for these proteins in innate defences in the absence of adaptive immunity, assessment of Lrrk2-linked and α-synuclein-based anti-viral properties in adult mice will allow us to investigate their role(s) in a mature immune system. In currently ongoing studies of testing additional RNA viruses in adult mice, we are examining their functions in the context of multiple factors, such as age as a determinant of disease progression, the influence of sex hormones, development beyond puberty, as well as responses via mature, adaptive immunity. In addition to this, testing of DNA-based pathogens (including those with double-stranded *vs*. single-stranded genomes) in the context of *Snca* and *Lrrk2* mutants will be warranted.

Taken together with our previous publications using the same virus (strain, dose) and same animal models (Shutinoski et al., 2019; Tomlinson et al., 2017), this study suggests that two PD-associated genes, *Lrrk2* and *Snca*, play a role in host defense against reovirus T3D-induced encephalitis primarily in the periphery rather than in brain-restricted host responses. These findings contribute to the increasing body of evidence indicating that both Lrrk2 and α-synuclein may have physiological roles outside of the brain, including those that may contribute to overall nervous system health through systemic immunity. Hence, further modeling of the numerous and complex interactions between environmental exposures and genetic risk factors in peripheral organs appears warranted when studying the pathogenesis of late-onset, neurodegenerative disorders, such as PD.

## Acknowledgment

This research was funded by the Parkinson Research Consortium Ottawa (M.O.L.; C.R), the Canadian Institutes of Health Research (M.G.S.; J.J.T.), Department of Medicine at The Ottawa Hospital (M.G.S.), and the Bhargava Family Research Chair in Neurodegeneration (M.G.S.). We are grateful to Dr. D. Shimshek and colleagues for sharing their *Lrrk2* mutant mouse lines.

## Author contributions

*Study design:* M.O.L., C.R., J.J.T., E.G.B., M.G.S.; *Writing and Figure preparation:* M.O.L., C.R., J.J.T., E.G.B., and M.G.S., who prepared drafts of the manuscript and figures. All authors reviewed and / or edited the manuscript and approved of the submitted version. *Experiments:* M.O.L. and C.R performed experiments and / or data analysis. *Analysis:* M.O.L., C.R., J.J.T., E.G.B., M.G.S. performed data interpretation. *Study supervision*: J.J.T., E.G.B., M.G.S. *Overall responsibility:* M.G.S.

## Competing interests

The authors declare they do not have any competing interests.

## Additional information

## Data and materials availability

Original data associated with this study are available in the main text and supplementary figures; additional data will be made available upon request.

## Correspondence and requests for materials

These should be addressed to J.J.T or M.G.S.

## Supplemental Figure 1

**Supplementary Fig. 1:**
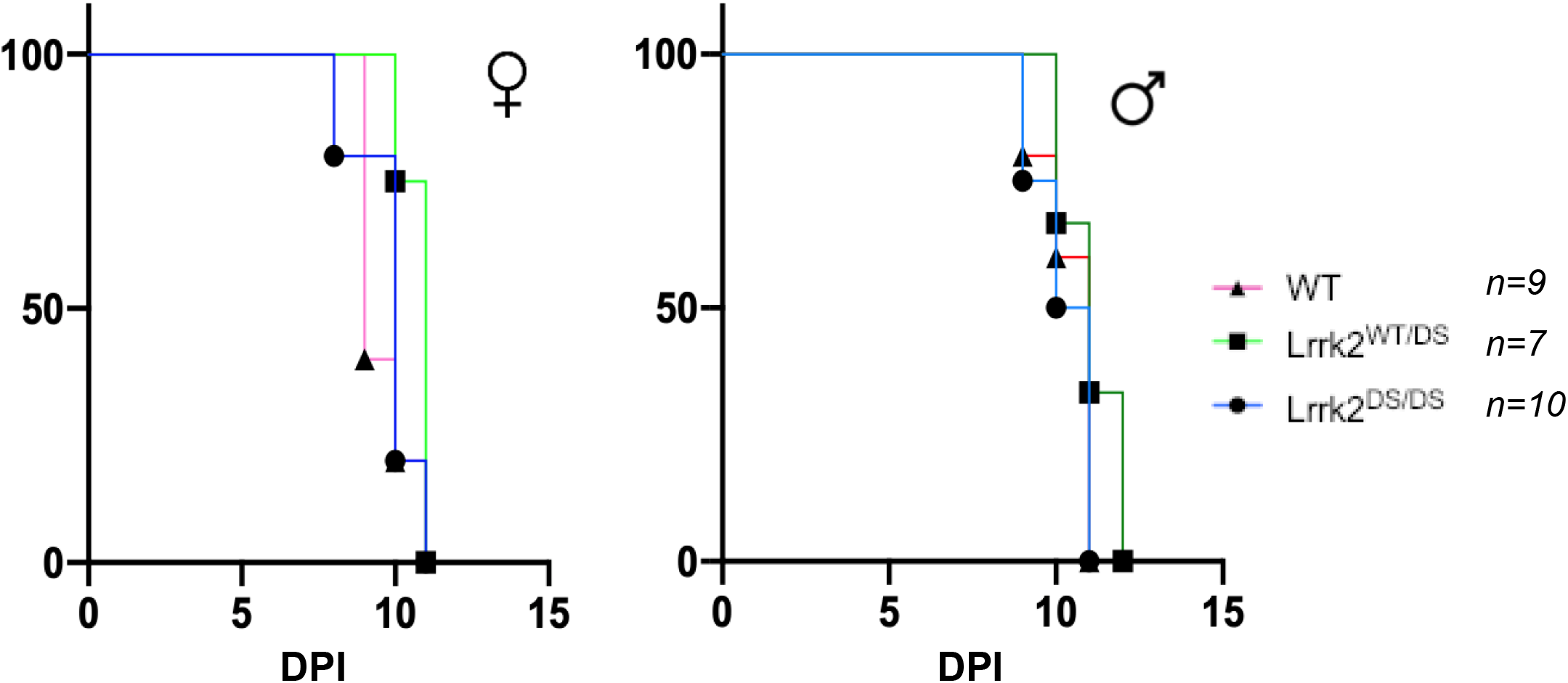
Intracerebral Inoculation with Reovirus T3D at Post-Natal Day 3 Does Not Uncover Genotypic Differences in Time-To-Death Between Wild-Type, Heterozygous, or Homozygous Lrrk2 p.D1994S Mutant Mice. Heterozygous Lrrk2 p.D1994S-carrying breeders were paired. At post-natal day 3 mice were inoculated with a dose of 5×10^2^ PFU of reovirus T3D via direct-brain injection into the left hemisphere and monitored for time-to-death. Inoculated pups were monitored over 14 days for a pre-determined moribund state. Time-to-death is graphically displayed as percent survival against days post inoculation (DPI), separated based on sex **(a)**. The *n* value indicates the sample size of sexes combined. Survival curves were statistically analyzed using Mantel-Cox (Log-Rank) tests. No significant group differences were observed.

## Notes

### Competing Interest Statement

The authors have declared no competing interest.

## References

Alam, M. M., Yang, D., Li, X.-Q., Liu, J., Back, T. C., Trivett, A., Karim, B., Barbut, D., Zasloff, M., & Oppenheim, J. J. (2022). Alpha synuclein, the culprit in Parkinson disease, is required for normal immune function. Cell Reports, 38(2), 110090. 10.1016/j.celrep.2021.110090

Bantle, C. M., Phillips, A. T., Smeyne, R. J., Rocha, S. M., Olson, K. E., & Tjalkens, R. B. (2019). Infection with mosquito-borne alphavirus induces selective loss of dopaminergic neurons, neuroinflammation and widespread protein aggregation. Npj Parkinson’s Disease, 5(1), 20. 10.1038/s41531-019-0090-8

Beatman, E. L., Massey, A., Shives, K. D., Burrack, K. S., Chamanian, M., Morrison, T. E., & Beckham, J. D. (2016). Alpha-Synuclein Expression Restricts RNA Viral Infections in the Brain. Journal of Virology, 90(6), 2767–2782. 10.1128/JVI.02949-15

Beckham, J. D., Tuttle, K. D., & Tyler, K. L. (2010). Caspase-3 activation is required for reovirus-induced encephalitis in vivo. Journal of Neurovirology, 16(4), 306–317. 10.3109/13550284.2010.499890

Braak, H., Tredici, K. D., Rüb, U., de Vos, R. A. I., Jansen Steur, E. N. H., & Braak, E. (2003). Staging of brain pathology related to sporadic Parkinson’s disease. Neurobiology of Aging, 24(2), 197–211. 10.1016/S0197-4580(02)00065-9

Cabin, D. E., Shimazu, K., Murphy, D., Cole, N. B., Gottschalk, W., McIlwain, K. L., Orrison, B., Chen, A., Ellis, C. E., Paylor, R., Lu, B., & Nussbaum, R. L. (2002). Synaptic Vesicle Depletion Correlates with Attenuated Synaptic Responses to Prolonged Repetitive Stimulation in Mice Lacking α-Synuclein. The Journal of Neuroscience, 22(20), 8797–8807. 10.1523/JNEUROSCI.22-20-08797.2002

Chen, S. G., Stribinskis, V., Rane, M. J., Demuth, D. R., Gozal, E., Roberts, A. M., Jagadapillai, R., Liu, R., Choe, K., Shivakumar, B., Son, F., Jin, S., Kerber, R., Adame, A., Masliah, E., & Friedland, R. P. (2016). Exposure to the Functional Bacterial Amyloid Protein Curli Enhances Alpha-Synuclein Aggregation in Aged Fischer 344 Rats and Caenorhabditis elegans. Scientific Reports, 6(1), 34477. 10.1038/srep34477

Gauvin, L., Bennett, S., Liu, H., Hakimi, M., Schlossmacher, M., Majithia, J., & Brown, E. G. (2013). Respiratory infection of mice with mammalian reoviruses causes systemic infection with age and strain dependent pneumonia and encephalitis. Virology Journal, 10(1), 67. 10.1186/1743-422X-10-67

Härtlova, A., Herbst, S., Peltier, J., Rodgers, A., BilkeilJGorzo, O., Fearns, A., Dill, B. D., Lee, H., Flynn, R., Cowley, S. A., Davies, P., Lewis, P. A., Ganley, I. G., Martinez, J., Alessi, D. R., Reith, A. D., Trost, M., & Gutierrez, M. G. (2018). LRRK2 is a negative regulator of Mycobacterium tuberculosis phagosome maturation in macrophages. The EMBO Journal, 37(12). 10.15252/embj.201798694

Herzig, M. C., Kolly, C., Persohn, E., Theil, D., Schweizer, T., Hafner, T., Stemmelen, C., Troxler, T. J., Schmid, P., Danner, S., Schnell, C. R., Mueller, M., Kinzel, B., Grevot, A., Bolognani, F., Stirn, M., Kuhn, R. R., Kaupmann, K., Van Der Putten, P. H., … Shimshek, D. R. (2011). LRRK2 protein levels are determined by kinase function and are crucial for kidney and lung homeostasis in mice. Human Molecular Genetics, 20(21), 4209–4223. 10.1093/hmg/ddr348

Jang, H., Boltz, D., Sturm-Ramirez, K., Shepherd, K. R., Jiao, Y., Webster, R., & Smeyne, R. J. (2009). Highly pathogenic H5N1 influenza virus can enter the central nervous system and induce neuroinflammation and neurodegeneration. Proceedings of the National Academy of Sciences, 106(33), 14063–14068. 10.1073/pnas.0900096106

Kalia, L. V., & Lang, A. E. (2015). Parkinson’s disease. The Lancet, 386(9996), 896–912. 10.1016/S0140-6736(14)61393-3

Kozina, E., Byrne, M., & Smeyne, R. J. (2022). Mutant LRRK2 in lymphocytes regulates neurodegeneration via IL-6 in an inflammatory model of Parkinson’s disease. Npj Parkinson’s Disease, 8(1), 24. 10.1038/s41531-022-00289-9

Leta, V., Urso, D., Batzu, L., Lau, Y. H., Mathew, D., Boura, I., Raeder, V., Falup-Pecurariu, C., van Wamelen, D., & Ray Chaudhuri, K. (2022). Viruses, parkinsonism and Parkinson’s disease: The past, present and future. Journal of Neural Transmission, 129(9), 1119–1132. 10.1007/s00702-022-02536-y

Liu, W., Liu, X., Li, Y., Zhao, J., Liu, Z., Hu, Z., Wang, Y., Yao, Y., Miller, A. W., Su, B., Cookson, M. R., Li, X., & Kang, Z. (2017). LRRK2 promotes the activation of NLRC4 inflammasome during Salmonella Typhimurium infection. Journal of Experimental Medicine, 214(10), 3051–3066. 10.1084/jem.20170014

Monogue, B., Chen, Y., Sparks, H., Behbehani, R., Chai, A., Rajic, A. J., Massey, A., Kleinschmidt-Demasters, B. K., Vermeren, M., Kunath, T., & Beckham, J. D. (2022). Alphasynuclein supports type 1 interferon signalling in neurons and brain tissue. Brain, 145(10), 3622–3636. 10.1093/brain/awac192

Passini, M. A., & Wolfe, J. H. (2001). Widespread Gene Delivery and Structure-Specific Patterns of Expression in the Brain after Intraventricular Injections of Neonatal Mice with an Adeno-Associated Virus Vector. Journal of Virology, 75(24), 12382–12392. 10.1128/JVI.75.24.12382-12392.2001

Peelaerts, W., Mercado, G., George, S., Villumsen, M., Kasen, A., Aguileta, M., Linstow, C., Sutter, A. B., Kuhn, E., Stetzik, L., Sheridan, R., Bergkvist, L., Meyerdirk, L., Lindqvist, A., Gavis, M. L. E., Van Den Haute, C., Hultgren, S. J., Baekelandt, V., Pospisilik, J. A., … Brundin, P. (2023). Urinary tract infections trigger synucleinopathy via the innate immune response. Acta Neuropathologica, 145(5), 541–559. 10.1007/s00401-023-02562-4

Shutinoski, B., Hakimi, M., Harmsen, I. E., Lunn, M., Rocha, J., Lengacher, N., Zhou, Y. Y., Khan, J., Nguyen, A., Hake-Volling, Q., El-Kodsi, D., Li, J., Alikashani, A., Beauchamp, C., Majithia, J., Coombs, K., Shimshek, D., Marcogliese, P. C., Park, D. S., … Schlossmacher, M. G. (2019). Lrrk2 alleles modulate inflammation during microbial infection of mice in a sex-dependent manner. Science Translational Medicine, 11(511). 10.1126/scitranslmed.aas9292

Spriggs, D. R., & Fields, B. N. (1982). Attenuated reovirus type 3 strains generated by selection of haemagglutinin antigenic variants. Nature, 297(5861), 68–70. 10.1038/297068a0

Tomlinson, J. J., Shutinoski, B., Dong, L., Meng, F., Elleithy, D., Lengacher, N. A., Nguyen, A. P., Cron, G. O., Jiang, Q., Roberson, E. D., Nussbaum, R. L., Majbour, N. K., El-Agnaf, O. M., Bennett, S. A., Lagace, D. C., Woulfe, J. M., Sad, S., Brown, E. G., & Schlossmacher, M. G. (2017). Holocranohistochemistry enables the visualization of α-synuclein expression in the murine olfactory system and discovery of its systemic anti-microbial effects. Journal of Neural Transmission, 124(6), 721–738. 10.1007/s00702-017-1726-7

Virgin, H. W., Bassel-Duby, R., Fields, B. N., & Tyler, K. L. (1988). Antibody protects against lethal infection with the neurally spreading reovirus type 3 (Dearing). Journal of Virology, 62(12), 4594–4604. 10.1128/jvi.62.12.4594-4604.1988

Zhang, Q., Pan, Y., Yan, R., Zeng, B., Wang, H., Zhang, X., Li, W., Wei, H., & Liu, Z. (2015). Commensal bacteria direct selective cargo sorting to promote symbiosis. Nature Immunology, 16(9), 918–926. 10.1038/ni.3233

