## Supplemental Figure 1 for "Variants in *Lrrk2* and *Snca* deficiency do not alter the course of primary encephalitis due to neurotropic reovirus T3D in newborn mice"

### Slide 1
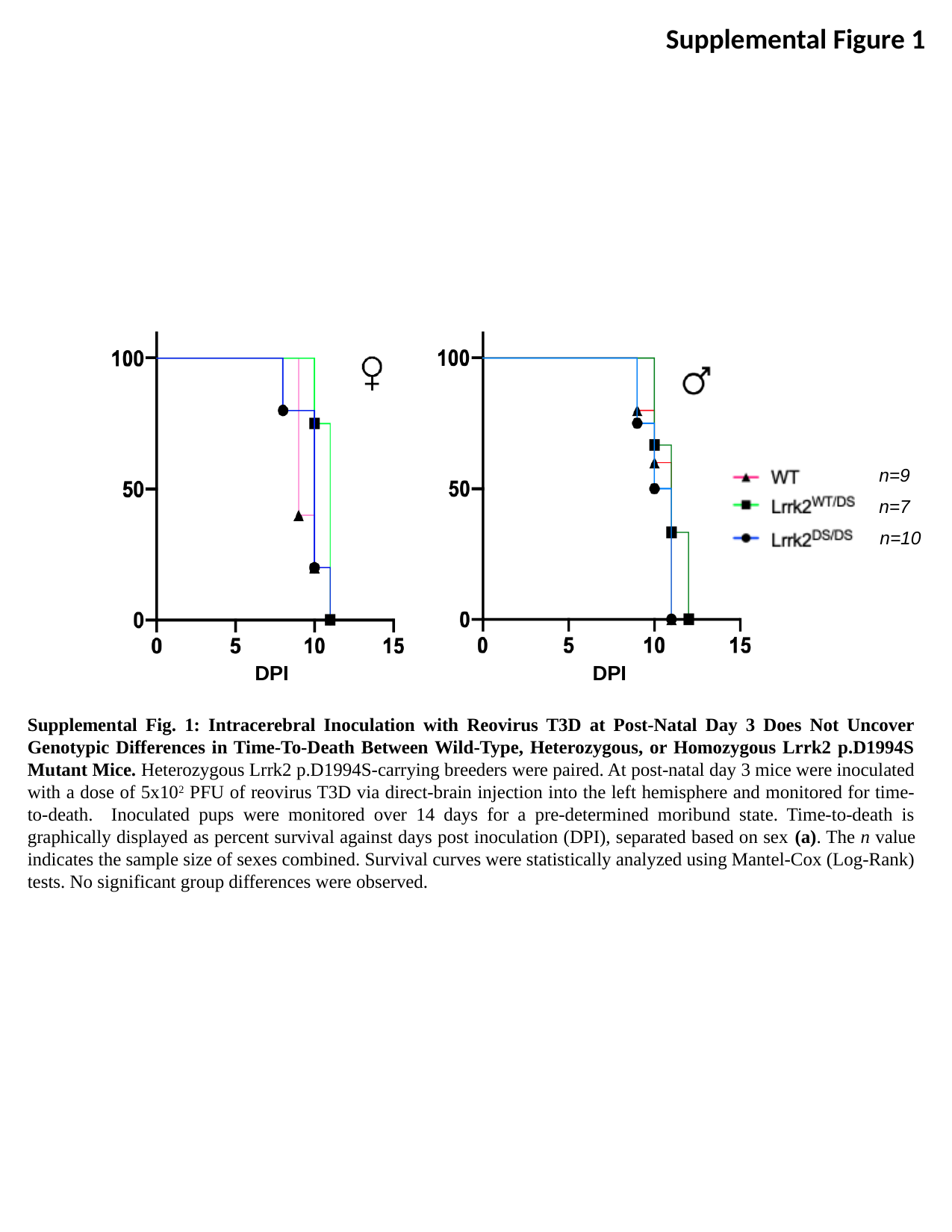

Supplemental Figure 1
n=9
n=7
n=10
DPI
DPI
Supplemental Fig. 1: Intracerebral Inoculation with Reovirus T3D at Post-Natal Day 3 Does Not Uncover Genotypic Differences in Time-To-Death Between Wild-Type, Heterozygous, or Homozygous Lrrk2 p.D1994S Mutant Mice. Heterozygous Lrrk2 p.D1994S-carrying breeders were paired. At post-natal day 3 mice were inoculated with a dose of 5x10^2^ PFU of reovirus T3D via direct-brain injection into the left hemisphere and monitored for time-to-death. Inoculated pups were monitored over 14 days for a pre-determined moribund state. Time-to-death is graphically displayed as percent survival against days post inoculation (DPI), separated based on sex (a). The n value indicates the sample size of sexes combined. Survival curves were statistically analyzed using Mantel-Cox (Log-Rank) tests. No significant group differences were observed.
